# Cryo-electron tomography reveals paracellular claudin-15 pores at the tight junction

**DOI:** 10.64898/2026.02.19.706806

**Authors:** Evgeniya A. Demchenko, Sarah McGuinness, Shaun Wood, Joseph Kainov, Victoria Pappas, Jotham Austin, Dominic Hyatt, Fatemeh Khalili-Araghi, Le Shen, Christopher R. Weber

**Affiliations:** Department of Biochemistry and Molecular Biology, The University of Chicago, Chicago, Illinois; Department of Biomedical Engineering, University of Illinois Chicago, Chicago, Illinois; Department of Physics, University of Illinois Chicago, Chicago, Illinois; Department of Pathology, The University of Chicago, Chicago, Illinois; Advanced Electron Microscopy Core, The University of Chicago, Chicago, Illinois; Department of Surgery, The University of Chicago, Chicago, Illinois

## Abstract

Tight junctions (TJs) are composed of anastomosing strands between epithelial cells. Members of the claudin family of proteins reside within TJ strands and either seal the paracellular space or assemble into charge and size-selective pathways. Functional studies suggest that claudin-mediated conductance pathways resemble traditional ion channels. However, such postulated pores have not been directly visualized. Using a model claudin deficient epithelium where exogenously introduced EGFP-CLDN15 is the only claudin family member expressed, our investigation sheds light on the arrangement and structure of the postulated claudin pores. Following correlative light and electron microscopical identification of TJs and cryo-electron tomography, we identified series of linearly distributed electron lucent features that locate between two closely apposed plasma membranes of adjacent cells. At these sites, the median spacing between adjacent features is 2.25 nm (IQR = 1.83), with a median 1.66 nm (IQR = 0.92) diameter. In contrast, such features were not observed in claudin deficient model epithelium with exogenous mCherry-ZO-1 expression. These findings agree with the postulated and extensively modeled claudin pores formed within the simple columnar epithelium. This provides the first direct evidence of paracellular pore organization and paves way for future biophysical investigation.

**SIGNIFICANCE:** By combining correlative fluorescence imaging, FIB milling, and cryo-ET within an epithelial system restricted to a single claudin isoform, we were able to visualize repetitive, low-density pore features within CLDN15-containing tight junctions (TJs), structures not previously resolved in intact epithelia. These features were absent in claudin-negative controls and displayed placement and geometry consistent with CLDN15 X-ray crystallography and molecular dynamics models. Quantitative measurements of pore diameter, paracellular gap width, and pore spacing further support their assignment as CLDN15 pores. These findings establish a structurally validated platform for defining claudin pore ultrastructure and provide a foundation for future efforts to compare pore-forming and barrier-forming claudins, understand disease-associated junction remodeling, and guide therapeutic modulation of epithelial barrier function.

## INTRODUCTION

Tight junctions (TJs) are located at the most apical end of the paracellular space that controls paracellular permeability(1, 2). They contain a branched network of anastomosing strands that has been visualized using freeze-fracture electron microscopy (3, 4). The claudin family of tetraspan membrane proteins comprises 27 mammalian members, and a number of these proteins can form TJ strands. Individual claudins either function primarily as barrier-forming proteins (e.g., CLDN1, 3, 4, 5) or as selectively permeable, charge- and size-selective paracellular ion pathways (e.g., CLDN2, 10b, 15) that regulate epithelial permeability (5–7). Claudin controlled charge- and size-selective permeability has been hypothesized to be due to ion channel pores, supported by our finding that CLDN2 conductance measured by TJ patch clamp technique demonstrates discrete single-channel opening and closing events (8).

Our understanding of 3-dimensional organization of claudin multimers has largely relied on crystal structures of linearly organized CLDN15 protomers, but these structures lack complete extracellular loops and do not capture interactions with claudins from adjacent cells (9). The partial CLDN15 crystal structure informed the earliest claudin-channel model (10), and was subsequently refined through molecular dynamics simulations (11–14). These studies converged on a model in which CLDN15 monomers assemble into opposing double rows that generate eight-subunit paracellular ion channels. However, these proposed claudin pores have not been directly visualized, leaving channel models incompletely validated. To overcome this limitation, we applied cryo-electron tomography (cryo-ET), which can resolve macromolecular assemblies in intact cells, to directly visualize CLDN15-dependent features within TJ strands.

## MATERIALS AND METHODS

A doxycycline-inducible CLDN15 expressing Madin Darby canine kidney (MDCK) II claudin quintuple-knockout (quinKO) (15) cell line was generated using PiggyBac-mediated stable integration and maintained under standard culture conditions. MDCK II quinKO monolayers with or without induced CLDN15 expression were used for immunofluorescence imaging, electrophysiologic measurements of barrier function, and cryo-electron tomography (cryo-ET). Transepithelial resistance, dilution potentials, and ion permeability measurements were obtained under current-clamp conditions using established methods.

For cryo-ET, cells were seeded onto collagen-coated EM grids, vitrified by plunge freezing (Leica, EM GP2 Automatic Plunge Freezer), and processed by correlative light and electron microscopy (cryo-CLEM) followed by cryo Ga-focused ion beam milling (Thermo Fisher Scientific, Aquilos 2 cryo-DualBeam) to generate 200–220-nm lamellae. Tilt series were acquired on a Thermo Fisher Scientific Titan Krios G3i microscope equipped with a Gatan K3 direct electron detector in CDS mode and BioQuantum energy filter, using a dose-symmetric ±54^°^ tilt scheme. Motion correction, tilt-series alignment, tomogram reconstruction, segmentation, and visualization was performed in IMOD (19). Model fitting and visualization completed in ChimeraX. Quantitative analyses of paracellular spacing, pore-to-pore distances, pore diameter, and membrane thickness were performed using FIJI and plotted with Python 3.9. Adobe Illustrator and Cinema4D were used for visualization. A complete description of plasmids, cell culture conditions, staining protocols, electrophysiology, vitrification parameters, cryo-CLEM and cryo-FIB/SEM workflows, tomogram reconstruction, segmentation, model alignment, and quantitative analysis methods is provided in the Supplemental Material.

## RESULTS AND DISCUSSION

### Generation and validation of MDCK II quinKO cells with induced EGFP-CLDN15 expression

Wild type epithelial cells express multiple species of pore forming and barrier forming claudins, which could lead to heterogenous protein organization, complicating morphological measurements. To circumvent this, we used a model epithelial system where only one dominant claudin species is expressed. This was achieved by using MDCK II quinKO line, in which genes for five endogenously expressed claudins are deleted, including CLDN1, CLDN2, CLDN3, CLDN4, and CLDN7 (15). Although RNA sequencing detected transcripts for additional claudins, including CLDN12 and CLDN16, their protein levels are very low and not concentrated at the TJ in MDCK II quinKO cells (16, 17). On this background, we introduced a previously validated tetracycline-inducible expression vector containing EGFP–CLDN15 expression cassette, providing a well-controlled system for studying CLDN15-dependent features. Upon doxycycline induction in confluent MDCK II quinKO-EGFP-CLDN15 monolayers, CLDN15 expression was verified by immunofluorescence, demonstrating that EGFP-CLDN15 colocalized with occludin at cell-cell junctions (FigureS1). To facilitate identification of TJ domains in MDCK II quinKO cells, we also generated a doxycycline-inducible mCherry–ZO-1 expressing line which was used to delineate TJ regions in selected experiments. Similar to previous reports using claudin knockout cell lines (17, 18), EGFP-CLDN15 expression significantly increased transepithelial resistance (FigureS2A) and enhanced Na^+^ selectivity (PNa/PCl) relative to non-induced monolayers or monolayers induced to express mCherry-ZO-1 (Figure S2B). This indicates EGFP-CLDN15 expression increases barrier function and Na^+^ selectivity.

### CLDN15 reorganizes plasma membranes of adjacent cells at tight junction-like domains

To visualize TJ strand organization, we performed cryo-electron tomography (cryo-ET) studies. Cryo-ET was chosen because it preserves subcellular organization in a near native state within intact cells (20, 21). MDCK II quinKO monolayers were grown directly on EM grids and induced to express either EGFP-CLDN15 or mCherry-ZO-1. Following vitrification, localization of TJ-like domains was determined by correlative light-electron microscopy (CLEM), and grids were milled at these sites. Subsequent to low-magnification cryo-EM mapping of a milled region, full resolution images were collected at these cell-cell contact sites as tilt series (FigureS3).

We first determined how EGFP-CLDN15 expression affects TJ-like membrane domain organization. Following tomogram reconstructions, adjacent cell membranes were segmented and meshed to produce 3D models (Figure 1A-F, Supplemental Movies 1-6). In both cell lines, sites with closely apposed plasma membranes are present, give multiple groove-like appearances within reconstructed plasma membranes, when viewed from inside of the cells. Tracking these grooves through the tilt series revealed frequent bending, and anastomoses of these grooves (black dashed lines in Figure 1A–F). To determine if EGFP-CLDN15 expression in MDCK II quinKO cells affects the distance between plasma membranes of adjacent cells at the grooved TJ-like region, the 50 smallest intermembrane distances in 3D per tomogram were measured directly from 3D-segmented cellular membranes. Tomogram-level median minimal intermembrane distances were reduced upon CLDN15 expression (two-sided Mann–Whitney U test, U = 30.0, p = 0.038). QuinKO tomograms exhibited a median distance of 5.88 nm (IQR = 0.27 nm, n = 3), whereas quinKO + CLDN15 tomograms showed a median distance of 4.57 nm (IQR = 1.18 nm, n = 11), (Figure 1G). This confirms that CLDN15 expression results in narrowing the paracellular space to bring opposing membranes closer and facilitate TJ formation. Our quantitative analyses agree with prior estimations freeze-fracture and TEM analyses in MDCK II WT and claudin quinKO cells (15). Otani et al. estimated 6–7 nm spacing between adjacent membranes at the most apical junctions in claudin quinKO cells, whereas we measure ∼6 nm. This difference likely reflects the more native preservation and volumetric sampling afforded by cryo-ET versus 2D TEM, where junction geometry can be skewed across sections. Furthermore, use of heavy metals during EM can produce variable binding or thickening in the imaging.

**Figure 1:**
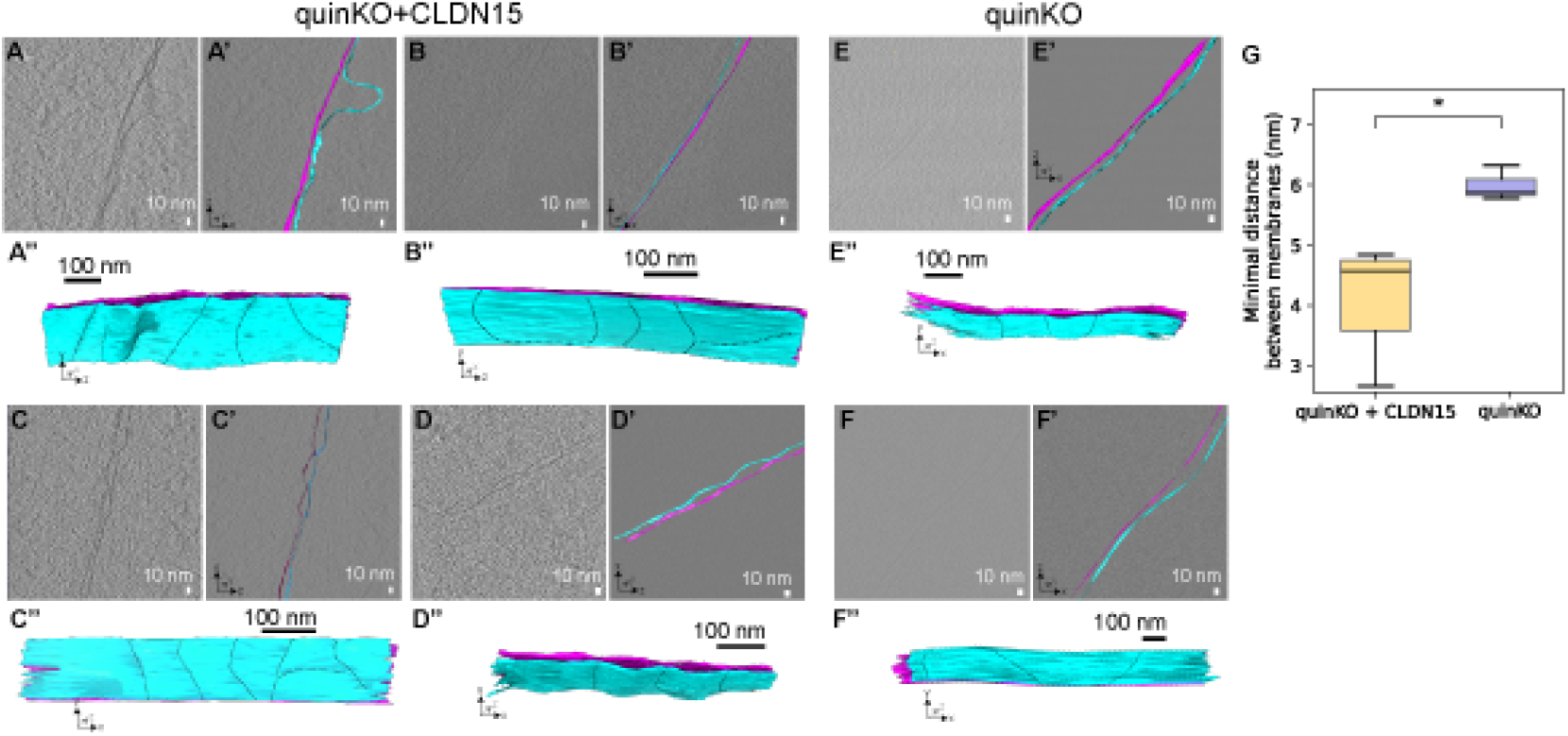
Claudin-15 expression reduces intermembrane spacing at TJ-like appositions in MDCK II quinKO cells. Representative reconstructed tomograms and 3D membrane models of MDCK II quinKO-EGFP-CLDN15 **(A-D)** and MDCK II quinKO-mCherry-ZO-1 **(E-F)** monolayers. Tomogram slices **(A-F)** with overlaid membrane segmentations **(A’-F’)** with opposing membranes in cyan and magenta) and **(A”-F”)** corresponding 3D models are shown (gray boxes indicate visible part of 3D model in tomogram slice). Black dashed lines indicate grooves observed on apposing cell membranes at TJ-like sites. **(G)** For each tomogram, 50 minimal three-dimensional intermembrane distances were extracted and median values were determined and shown. quinKO + CLDN15 showed a median distance of 4.57 nm (IQR = 1.18 nm, n = 11) and quinKO showed a median distance of 5.88 nm (IQR = 0.27 nm, n = 3). (mCherry–ZO-1, n = 3 tomograms and EGFP–CLDN15, n = 11 tomograms), * p < (two-sided Mann–Whitney U test, U = 30.0, p = 0.038).

### Visualization of claudin-15–mediated paracellular pores

We next examined the organization of these CLDN15-dependent TJ-like domains (represented schematically in Figure 2A) observed in tomogram slices (Figure 2B). Within these tomogram slices, multiple repetitive, linearly arranged low electron density foci were frequently observed at TJ domains (Figure 2C,C’, D,D’; yellow arrows). In contrast, such foci were not observed in MDCK II quinKO cells with mCherry-ZO-1 expression. We quantified membrane thickness, width of the paracellular space, pore diameter of claudin pores, and pore-to-pore distances for linear clusters of pores (Figure 3A). In tomogram snapshots of closely apposed plasma membranes between adjacent cells, the median plasma membrane thickness is 4.36 nm (IQR=0.95, n=22) (Figure 3B), and the median width of the paracellular space at these sites was 4.33 nm (IQR=0.73, n=11) (Figure 3C). The median inter-membrane distance of 4.57 nm for CLDN15-expressing monolayers (Figure 1G) is consistent with the median paracellular space width (4.33 nm) collected from 2D tomogram slices (Figure 3C). Previously published all-atom molecular dynamics (11, 12) report that the pore’s *in silico* mean diameter at the widest part of the CLDN15 pore is ∼1.6 nm, consistent with the median 1.55 nm (IQR = 0.92, n = 41), diameter of the observed features *in situ* (Figure 3D). From tomogram snapshots, distances between individual CLDN15 pores arranged in linear clusters have been measured, having a median feature-to-feature spacing of 2.25 nm (IQR=1.83, n=77) (Figure 3E). We report the median distance between CLDN15 pores based on our molecular-dynamics–derived computational model of a 8-pore CLDN15 strand (11) as (Figure 3A) 3.18 nm (IQR=0.21, n=7) and the 3-pore model of CLDN15 (12) revealed a periodicity between CLDN15 monomers with median 2.83 nm (IQR=0.09, n=2) (Figure 3E). Similarities between our quantitative measurements of feature to feature spacing in this work and quantitative measurements of previously published computational models (11, 12) suggest that the structures indicated are CLDN15 pores.

**Figure 2:**
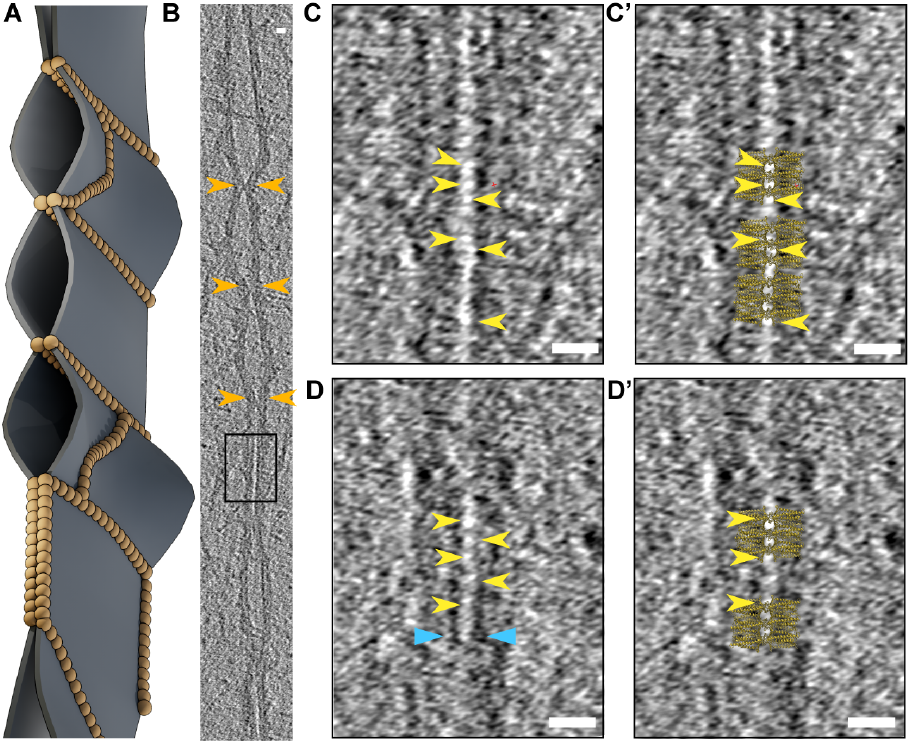
Claudin-15 forms low electron density foci at tight junctions-like domains. **(A)** 3D schematic of claudin strands (gold) depicted as beads-on-a-string. **(B)** Tomogram snapshot of MDCK II quinKO cells with EGFP-CLDN15 showing membrane “kissing points” (gold arrows). Prolonged membrane appositions are also evident (box). **(C)** Zoomed view of the boxed region in B, revealing linearly arranged paracellular low electron density foci (yellow arrows). **(D)** Representative tomogram slice from a different TJ-like domain, showing similar foci (yellow arrows) between the upper leaflets of closely apposed plasma membranes (blue arrows) **(C’, D’)** Overlay of three-pore computational model of CLDN15 (12) onto tomogram snapshots (Scale bar = 10 nm).

**Figure 3:**
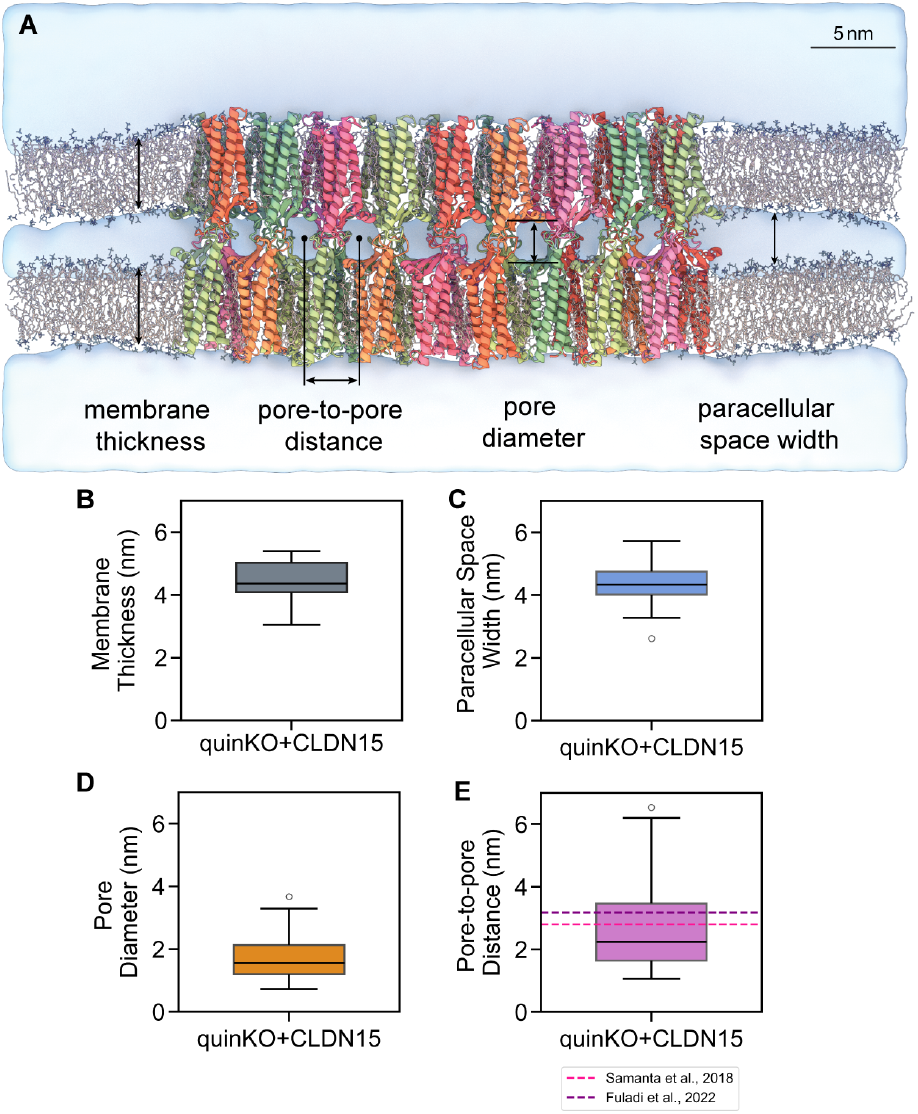
Quantitative analysis of claudin-15 dependent low electon density foci. **(A)** Computational model of 8-pore, 36-monomer CLDN15 strand embedded in apposing plasma membranes with schematic indicating measurments of membrane thickness, pore-to-pore distance, pore diameter, and paracellular space width (distance between two closely apposed plasma membranes). **(B)** Median plasma membrane thickness is 4.36 nm (IQR=0.95, n=22). **(C)** Median width of the paracellular space is 4.33 nm (IQR=0.73, n=11).**(D)** Diameters of low electron density foci between opposing plasma membranes have a median value of 1.55 nm (IQR = 0.92, n = 41). **(E)** Center-to-center spacing between low electron densities has a median of 2.25 nm (IQR=1.83, n=77). Dotted lines indicate the pore-to-pore distance measured in molecular dynamics simulations reported in prior studies: 3-pore model has median of 2.83 nm (IQR=0.09, n=2) (12) and 8-pore model median is 3.18 nm (IQR=0.21, n=7) (11)

## CONCLUSION

Based on morphological and quantitative features, such as low electron density, periodicity, spatial confinement within CLDN15 positive junctions confirmed by CLEM, and known ultrastructural features of TJ strand organization, we have identified CLDN15 pores in our cryo-ET data sets. The small paracellular width measurements support the conclusion that CLDN15 defines the tightest intermembrane appositions and promotes TJ formation. Although there is good geometric agreement on average between experimentally determined spacing of TJ pores and computational models, the values are not identical. In our experimental data, distances between these loci are not always uniform. We observed loci that are positioned more closely together, appearing as doublets, and others that are farther apart. We attribute these variations to several factors. First, reconstructed sections of tomograms may not capture each pore at the same axial position, as computationally determined TJ pores have wide and narrow regions. Second, cryo-ET may capture pores at different stages of assembly or disassembly. Third, CLDN15 distribution along TJ strands may be discontinuous. Fourth, TJ strands exhibit intrinsic structural heterogeneity, including regions of fusion, bending, and curvature, that can reduce the number of pores present within a given tomographic plane. Other, undefined mechanisms may also contribute. Additional studies are needed to define these differences. Notably, direct visualization of pore-like structures was rare, with clear pores resolved in only 4 of 38 CLDN15-positive TJ sites. This is expected given geometric and technical constraints of cryo-ET (22, 23), including variability in membrane orientation relative to the milling and imaging planes, overlapping densities within the lamella that can obscure small features, and ice-thickness variability. In addition, the small (2 nm) pore size approaches the effective resolution limit of 200-nm lamellae imaged over a ±54^°^ tilt range, making it difficult to acquire datasets large enough and sufficiently consistent for sub-tomogram averaging. Consequently, infrequent pore detection and deviations from idealized spacing primarily reflect sampling and projection limitations rather than the absence of CLDN15 pores.

This proof-of-concept data opens doors for understanding TJ structure and function with unprecedented detail. Future studies will build on this foundation by generating expanded datasets suitable for sub-tomogram averaging, enabling higher-resolution visualization of claudin pore architecture and more definitive identification of structural classes. Beyond CLDN15, this approach will be extended to define the similarities and differences among structures formed by individual pore-forming and barrier-forming claudins, as well as to determine how mixtures of claudin species within the same TJ strand collectively shape claudin pore and strand organization, and permeability. Because TJs exhibit substantial spatial heterogeneity, including differences along cell-to-cell contacts, at tricellular junctions, within curved membrane regions, and at sites of strand fusion or branching, and because their architecture varies across tissues and species, future work will map these contextual variations to determine how membrane geometry and local regulatory cues influence pore organization. Finally, studies using cell lines, organoids and tissues will allow us to define how disease and inflammation remodel claudin organization and pore structure, linking ultrastructural changes to barrier dysfunction in conditions such as inflammatory bowel disease, infectious colitis, and pancreatitis. Together, these efforts will provide a more complete biophysical understanding of TJ organization and claudin interactions, ultimately enabling the rational development of small-molecule, peptide, and antibody-based modulators of TJ function for therapeutic applications.

## AUTHOR CONTRIBUTIONS

CRW, LS, FKH, EAD, SM, JA conceptualized the project and designed experiments. SW and JK generated cell lines, SW performed line characterization experiments. EAD managed cell culture and prepared cryo-grids. EAD, VP, JA performed EM data collection. SM, VP, JA reconstructed tomograms. EAD, SM, DH built 3D segmentations. SM, EAD carried out Cryo-ET data analysis and visualization. All authors contributed to the writing of the manuscript.

## ACKNOWLEDGMENTS

We are thankful to Dr. Eduardo Perozo for discussions and support throughout the project; the University of Chicago Advanced Electron Microscopy Facility (RRID:SCR019198) for microscopes maintenance and training. This work was supported by a Chicago Biomedical Consortium catalyst award (C-106), NIH R01DK131542 and NSF MCB-1846021.

## DECLARATION OF INTERESTS

The authors declare no competing interests.

## SUPPLEMENTAL INFORMATION

### SUPPLEMENTAL METHODS

#### Molecular cloning, cell culture, and stable cell line generation

Madin-Darby canine kidney (MDCK) II line (15) with CLDN1, CLDN2, CLDN3, CLDN4, CLDN7 knockout (quintuple KO, quinKO) was used to generate stable cell lines used for this study. Doxycycline-inducible EGFP-human CLDN15 or mCherry-ZO1 expressing PiggyBac constructs were stably transfected into MDCK II quinKO cells together with a PiggyBac transposase encoding plasmid (24). Following hygromycin selection, pooled clones were used for subsequent experiments. Cells were maintained in low-glucose Dulbecco’s Eagle Modified Medium (DMEM, Gibco) supplemented with 10% fetal bovine serum (FBS, Gibco), 15mM HEPES, and 100 mg/ml hygromycin, and incubated at 37^°^C in a 5% CO2-containing atmosphere.

**Figure S1:**
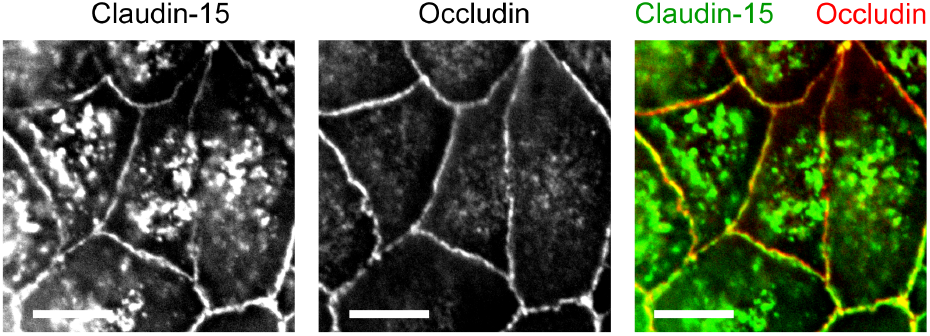
Generation of tet-inducible EGFP-CLDN15 - expressing MDCK II quinKO monolayers. After doxycycline treatment, fluorescent microscopy showed induced EGFP-CLDN15 expression (green) is detected at cell–cell junctions and co-localizes with antibody stained endogenous occludin (red), a transmembrane tight junction protein. (Scale bar=10 *µ*m)

#### Fluorescent staining and microscopy

Confluent MDCK II quinKO cell monolayers with or without doxycycline induced CLDN15 or ZO-1expression cultured on permeable supports were fixed in 1% paraformaldehyde prepared in phosphate-buffered saline (PBS, pH 7.4) for 1hr at room temperature. Following quenching in 50 mM NH4Cl in PBS and PBS wash, monolayers were permeabilized and incubated with blocking buffer (0.05% saponin and 3% BSA in PBS) for 30 min. Primary antibodies against CLDN1, CLDN15, occludin, and ZO-1 (Thermo Fisher Scientific) were diluted in blocking buffer and incubated with monolayer at 4 ^°^C overnight. Following washes with blocking buffer, monolayers were incubated with blocking buffer supplemented with Alexa Fluor 488-or 594-conjugated secondary antibodies (Jackson Immunolabs) and Hoechst 33342(Thermo Fisher Scientific) for 1hr at room temperature. Following blocking buffer washing, monolayers were cut from semipermeable supports and mounted onto glass slides with ProLong Diamond mounting medium(Thermo Fisher Scientific). Super resolution imaging was performed using a Leica SP8 inverted laser scanning confocal microscope equipped with a white light laser and 3D STED (Leica Microsystems, Wetzlar, Germany). Images were collected with a 100× 1.4 NA HC PL APO CS2 oil immersion objective, and STED depletion lasers at 592 nm and 775 nm were used for enhanced lateral and axial resolution. Z-stacks were acquired at 0.1–0.2 *µ*m intervals, and deconvolution was performed using Leica LAS X software. Final images were analyzed in ImageJ (NIH).

**Figure S2:**
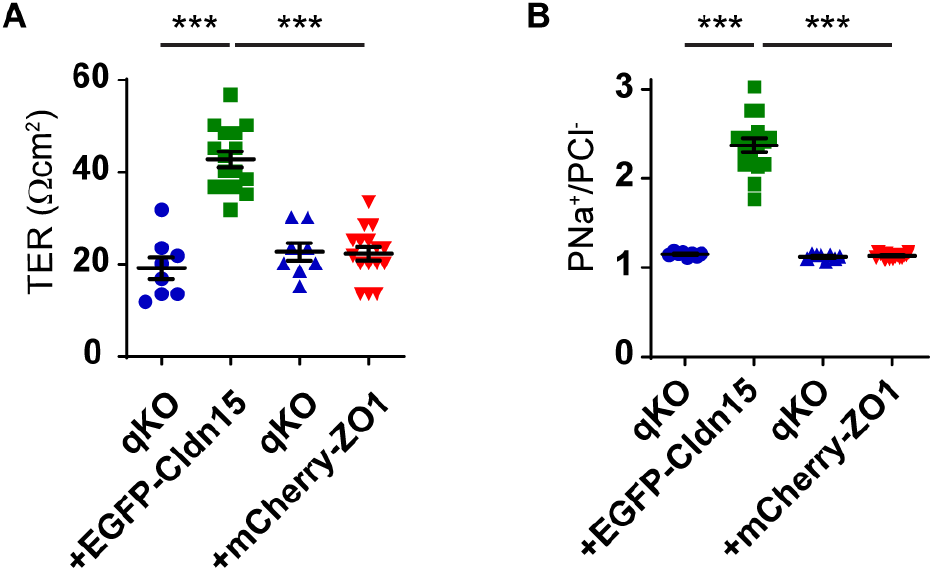
Electrophysiological effects of EGFP-CLDN15 expression on MDCK II quinKO monolayers. **(A)** Doxycycline-induced EGFP-CLDN15 expression (green squares) significantly increased transepithelial electrical resistance (TER) in MDCK II quinKO monolayers (blue circles, p<0.001, Welch’s one-way ANOVA with Dunnet’s post hoc test), while doxycycline-induced mCherry-ZO-1 expression (red down-pointing triangles) did not affect TER in a separate stable cell line (blue up-pointing triangles). **(B)** Doxycycline-induced EGFP-CLDN15 expression (green squares) significantly increased relative permeability between Na+ and Cl- (PNa+/PCl-) in MDCK II quinKO monolayers (blue circles, p*<*0.001, Welch’s one-way ANOVA with Dunnet’s post hoc test), suggesting increased Na+ permeability. In contrast, doxycycline-induced mCherry-ZO-1 expression (red down-pointing triangles) did not affect TER in a separate stable cell line (blue up-pointing triangles).

#### Electrophysiology

Barrier function of MDCKII quinKO monolayers with or without doxycycline treatment was evaluated under current-clamp conditions using a 558C-5 voltage-clamp system (University of Iowa, Iowa City, IA). Cells were grown on semipermeable Transwell inserts (0.4 *µ*m pore size; Corning) and bathed bilaterally with Hank’s balanced salt solution (HBSS, 138 mM NaCl, 0.3 mM Na_2_HPO_4_, 0.4 mM MgSO_4_·7H_2_O, 0.5 mM MgCl_2_·6H_2_O, 5.0 mM KCl, 0.3 mM KH_2_PO_4_, 15 mM HEPES, 1.3 mM CaCl_2_, and 5.5 mM glucose; pH 7.4). Ag/AgCl electrodes connected via 3 M KCl agar bridges were used to record transepithelial potential and current transepithelial electrical resistance (TER) was calculated. For dilution potential measurements, the basal chamber solution was replaced with HBSS with 50% NaCl, which was osmotically balanced with mannitol. The resulting potential, with transepithelial current clamped to zero, was used to calculate cation-to-anion permeability ratios (PNa^+^/PCl^™^) as described previously (25).

#### Preparation and vitrification of cell-seeded EM grids

C-flat Holey Carbon Grids (2/1, Au, 200 mesh) were plasma-cleaned for 30 s in an air mixture in a Solarus Plasma Cleaner (Gatan), then exposed to UV light in a tissue culture hood for 10 mins. Sterilized grids were coated with collagen, washed with PBS, and transferred to cell culture dish containing cell culture medium. MDCK II cells as described above were trypsinized, resuspended in culture medium, and seeded onto the grids dropwise at 20-40% confluency. The next day gene expression was induced with 25 *µ*g/mL doxycycline and incubated at 37^°^C in 5% CO2-containing atmosphere for 3-4 days before vitrification. The cells were frozen in liquid-nitrogen-cooled liquid ethane using an EM GP2 Automatic Plunge Freezer (Leica) with the following parameters: 100 % humidity, 22^°^ C chamber temperature, back-side blotting, and 25-70 s blot time depending on the cell density before freezing. Frozen grids were stored under liquid nitrogen. Live-cell imaging before freezing and vitrified grid overview is shown in Figure S3 A–C.

**Figure S3:**
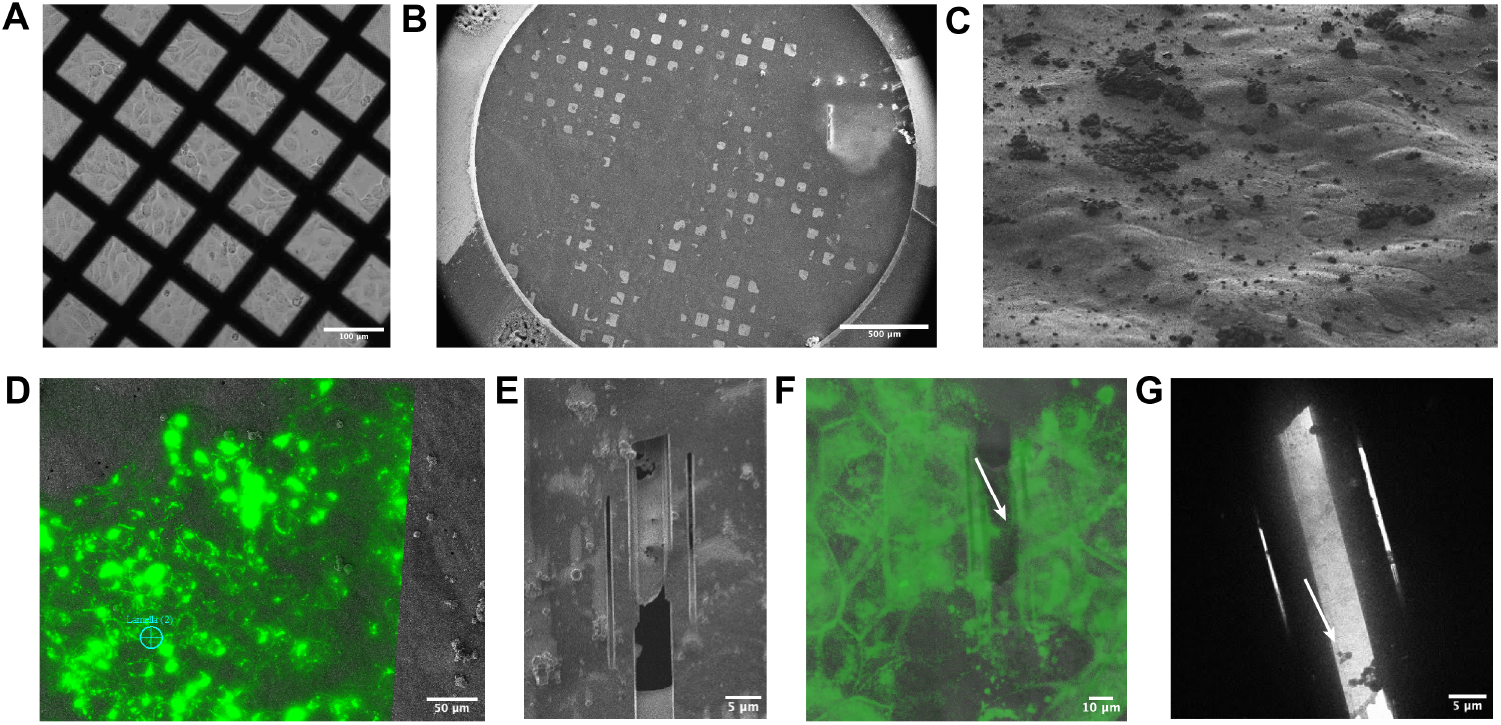
Sample preparation workflow for Cryo-ET of tight junctions. **(A)** Live cell imaging of EGFP-CLDN15–expressing MDCK II quinKO cells grown on EM grids (Scale bar = 100 *µ*m). **(B)** Overview of vitrified grid (Scale bar = 500 *µ*m). **(C)** Tilted view of grid square showing epithelial monolayer. **(D)** CLEM image used to define lamella position (blue cross; Scale bar 50 = *µ*m). **(E)** FIB-milled lamella (Scale bar 5 *µ*m). **(F)** Post-milling CLEM evaluation of the milled lamella. White arrow indicates an EGFP-CLDN15–positive cell-cell contacts (Scale bar 10 *µ*m). **(G)** Low magnification map of the lamella acquired on the Krios electron microscope; arrow indicates a tight junction (Scale bar = 5 *µ*m).

#### Cryo-CLEM and Cryo-FIB/SEM lamella preparation

Frozen grids were clipped into AutoGrid Rings (Thermo Fisher Scientific) with a cutout on one side for better access for the ion beam and loaded into an Aquilos 2 dual-beam instrument (Thermo Fisher Scientific). Samples were first sputter-coated in-column with Pt for 15 s at 10 Pa and 20 mA, followed by organic platinum deposition via a gas injection system for 20–30 seconds. To determine apical end of cell-cell contact site in MDCK II quinKO cells, cryo-CLEM (Correlative Light and Electron Microscopy) was performed using an iFLM detector (Thermo Fisher Scientific). Fluorescent signal of EGFP-CLDN15 and mCherry-ZO-1 was used to identify TJ-like sites between stably transfected MDCK II quinKO cells (Figure S3D). Correlation of the acquired EM and fluorescent images was performed in Maps software (Thermo Fisher Scientific) by aligning features visible in both reflection and EM images. Cells were Cryo-FIB milled at a 9^°^ target angle using AutoTEM software (Thermo Fisher Scientific). The automated protocol included generation of stress relief cuts at 1 nA, followed by sequential rough (0.3-0.5 nA), medium (0.1-0.3nA), and fine (50 pA-0.1nA) milling, concluded with two polishing steps at 30 and 10 pA respectively. Final lamellae had a target thickness of 200–220 nm. Milled lamellae were re-evaluated by fluorescence using the iFLM detector to confirm remaining target presence (Figure S3 E–F). Milled samples were stored under liquid nitrogen.

#### CryoET data collection

Cryo-ET data collection was performed on a Titan Krios G3i (Thermo Fisher Scientific) 300 kV transmission electron microscope using a K3 direct-electron detector in CDS mode and BioQuantum energy filter with a 20 eV slit (Gatan). Tilt series were collected with Tomography 5 (Thermo Fisher Scientific) at magnification of 53000 x and physical pixel size of 1.68 Å. Datasets were collected using dose-symmetric tilt scheme starting from the lamella pre-tilt angle (9^°^) with a 3^°^ tilt increment and angular range of ±54^°^. The accumulated dose of each tilt series was 110-120 e/Å^2^ with a defocus range of -4 to -6 *µ*m. 69 tilt series were obtained for MDCK II quinKO monolayers with EGFP-CLDN15 expression and 29 tilt series were obtained for MDCK II quinKO monolayers with mCherry-ZO-1 expression samples (Figure S3G).

#### Data processing, tomogram reconstruction, and model fitting

Motion correction, tilt-series alignment and tomogram reconstruction were completed utilizing IMOD software with eTomo graphic user interface (19). Tomograms were reconstructed using back-projection and 3D CTF correction. IMOD was also used for manual annotation and segmentation of tomograms (26). To segment the plasma membranes, contours were drawn manually every 10-50 frames, linearly interpolated and meshed with a thickness of 1. 14 tomograms were segmented, including 3 MDCK II quinKO monolayers with mCherry-ZO-1 expression and 11 MDCK II quinKO monolayers with EGFP-CLDN15 expression. Visualization and figure generation were completed in FIJI, UCSF ChimeraX, IMOD, Cinema4D, and Adobe Illustrator. A previously published computational model (11) of CLDN15 strands was used for alignment and visual comparison, performed in UCSF ChimeraX.

#### Pore measurements

Tomogram snapshots containing TJ pores were collected. Using FIJI, linear density profiles of areas containing TJ pores were extracted. Distances between adjacent pores were measured from pore center to pore center, which was also compared to the pore-to-pore center distance of our 8-pore computational model (simulated for 1 *µ*s). Linear density profiles of opposing plasma membranes were calculated. To manually measure the width of the paracellular space and the thickness of the two plasma membranes for quinKO and quinKO+CLDN15 groups, linear density profiles were utilized to define the edges of the plasma membranes. To measure pore size from linear density profiles, each pore is treated as an ellipse, and the length of the major axis was collected. Pore and adjacent membrane measurements were plotted as boxplots using Python3.

#### Quantification and statistical analysis

For each tomogram, all pairwise distances between opposing non interpolated 3D segmented plasma membranes were computed in three-dimensional space using a python script. From these distributions, the 50 smallest intermembrane distances were selected to enrich for measurements within TJ–like regions. To ensure statistical independence of biological replicates, each tomogram was treated as a single experimental unit. The median of the 50 minimal distances was calculated for each tomogram, yielding one representative value per tomogram. These per-tomogram medians were then used for group-level comparisons between conditions. Results with P values less than 0.05 were considered statistically significant. The boxplot was plotted using Python3.

## Notes

https://uofi.box.com/s/wd9444kd568l76v6csreclatgm1n2ftp

### Competing Interest Statement

The authors have declared no competing interest.

## REFERENCES

1. Shen, L., C. R. Weber, D. R. Raleigh, D. Yu, and J. R. Turner, 2011. Tight junction pore and leak pathways: a dynamic duo. Annual Review of Physiology 73:283–309.

2. Farquhar, M. G., and G. E. Palade, 1963. Junctional complexes in various epithelia. Journal of Cell Biology 17:375–412.

3. Claude, P., 1978. Morphological factors influencing transepithelial permeability: a model for the resistance of the zonula occludens. Journal of Membrane Biology 39:219–232.

4. Chalcroft, J. P., and S. Bullivant, 1970. An interpretation of liver cell membrane and junction structure based on observation of freeze-fracture replicas of both sides of the fracture. Journal of Cell Biology 47:49–60.

5. Furuse, M., K. Fujita, T. Hiiragi, K. Fujimoto, and S. Tsukita, 1998. Claudin-1 and -2: novel integral membrane proteins localizing at tight junctions with no sequence similarity to occludin. Journal of Cell Biology 141:1539–1550.

6. Furuse, M., H. Sasaki, and S. Tsukita, 1999. Manner of interaction of heterogeneous claudin species within and between tight junction strands. Journal of Cell Biology 147:891–903.

7. Günzel, D., and A. S. L. Yu, 2013. Claudins and the modulation of tight junction permeability. Physiological Reviews 93:525–569.

8. Weber, C. R., G. H. Liang, Y. Wang, S. Das, L. Shen, A. S. L. Yu, D. J. Nelson, and J. R. Turner, 2015. Claudin-2-dependent paracellular channels are dynamically gated. eLife 4:e09906.

9. Suzuki, H., T. Nishizawa, K. Tani, Y. Yamazaki, A. Tamura, R. Ishitani, N. Dohmae, S. Tsukita, O. Nureki, and Y. Fujiyoshi, 2014. Crystal structure of a claudin provides insight into the architecture of tight junctions. Science 344:304–307.

10. Suzuki, H., K. Tani, A. Tamura, S. Tsukita, and Y. Fujiyoshi, 2015. Model for the architecture of claudin-based paracellular ion channels through tight junctions. Journal of Molecular Biology 427:291–297.

11. Fuladi, S., S. McGuinness, L. Shen, C. R. Weber, and F. Khalili-Araghi, 2022. Molecular mechanism of claudin-15 strand flexibility: a computational study. Journal of General Physiology 154:e202213116.

12. Samanta, P., Y. Wang, S. Fuladi, J. Zou, Y. Li, L. Shen, C. Weber, and F. Khalili-Araghi, 2018. Molecular determination of claudin-15 organization and channel selectivity. Journal of General Physiology 150:949–968.

13. McGuinness, S., S. Sajjadi, C. R. Weber, and F. Khalili-Araghi, 2024. Computational models of claudin assembly in tight junctions and strand properties. International Journal of Molecular Sciences 25:3364.

14. Alberini, G., F. Benfenati, and L. Maragliano, 2018. Molecular dynamics simulations of ion selectivity in a claudin-15 paracellular channel. Journal of Physical Chemistry B 122:10783–10792.

15. Otani, T., T. P. Nguyen, S. Tokuda, K. Sugihara, T. Sugawara, K. Furuse, T. Miura, K. Ebnet, and M. Furuse, 2019. Claudins and JAM-A coordinately regulate tight junction formation and epithelial polarity. Journal of Cell Biology 218:3372–3396.

16. Pouyiourou, I., A. Fromm, J. Piontek, R. Rosenthal, M. Furuse, and D. Günzel, 2025. Ion permeability profiles of renal paracellular channel-forming claudins. Acta Physiologica 241:e14264.

17. Furuse, M., D. Nakatsu, W. Hempstock, S. Sugioka, N. Ishizuka, K. Furuse, T. Sugawara, Y. Fukazawa, and H. Hayashi, 2023. Reconstitution of functional tight junctions with individual claudin subtypes in epithelial cells. Cell Structure and Function 48:1–17.

18. Kashihara, H., H. Tanaka, M. Kitamata, G. Shiratsuchi, T. Katsuno, K. Tsukita, T. Nishida, M. Hamasaki, F. Eisenstein, H. Suzuki, S. Nakamura, K. Aoyama, T. Yagi, R. Danev, Y. Fujiyoshi, A. Tamura, and S. Tsukita, 2025. Functional landscape of mechanistic diversity in 27 claudin family members at tight junctions 11:eadx7431.

## SUPPLEMENTAL REFERENCES

19. Kremer, J. R., D. N. Mastronarde, and J. R. McIntosh, 1996. Computer visualization of three-dimensional image data using IMOD. Journal of Structural Biology 116:71–76.

20. Hong, Y., Y. Song, Z. Zhang, and S. Li, 2023. Cryo-electron tomography: the resolution revolution and a surge of in situ virological discoveries. Annual Review of Biophysics 52:339–360.

21. Hylton, R. K., and M. T. Swulius, 2021. Challenges and triumphs in cryo-electron tomography. iScience 24:102959.

22. Watson, A. J. I., and A. Bartesaghi, 2024. Advances in cryo-ET data processing: meeting the demands of visual proteomics. Current Opinion in Structural Biology 87:102861.

23. Turk, M., and W. Baumeister, 2020. The promise and the challenges of cryo-electron tomography. FEBS Letters 594:3243–3261.

24. Li, Z., I. P. Michael, D. Zhou, A. Nagy, and J. M. Rini, 2013. Simple piggyBac transposon-based mammalian cell expression system for inducible protein production. Proceedings of the National Academy of Sciences of the USA 110:5004–5009.

25. Weber, C. R., D. R. Raleigh, L. Su, L. Shen, E. A. Sullivan, Y. Wang, and J. R. Turner, 2010. Epithelial myosin light chain kinase activation induces mucosal interleukin-13 expression to alter tight junction ion selectivity. Journal of Biological Chemistry 285:12037–12046.

26. Danita, C., W. Chiu, and J. G. Galaz-Montoya, 2022. Efficient manual annotation of cryogenic electron tomograms using IMOD. STAR Protocols 3:101658.

